# Transmission of human influenza A virus in pigs selects for adaptive mutations on the HA gene

**DOI:** 10.1101/2022.04.04.487085

**Authors:** Jong-suk Mo, Eugenio J. Abente, Troy C. Sutton, Lucas M. Ferreri, Ginger Geiger, Phillip C. Gauger, Daniel R. Perez, Amy L. Vincent Baker, Daniela S. Rajao

## Abstract

Influenza A viruses (IAV) cause respiratory diseases in many host species, including humans and pigs. The spillover of IAV between swine and humans has been a concern for both public health and the swine industry. With the emergence of the triple reassortant internal gene (TRIG) constellation, establishment of human-origin IAVs in pigs has become more common, leading to increased viral diversity. However, little is known about the adaptation processes that are needed for a human-origin IAV to transmit and become established in pigs. We generated a reassortant IAV containing surface gene segments from a human IAV strain and internal gene segments from the 2009 pandemic and TRIG IAV lineages and demonstrated that it can replicate and transmit in pigs. Sequencing and variant calling analysis identified a mutant that emerged during replication in pigs, which was mapped to a region near the receptor binding site of the hemagglutinin (HA). The variant was present in all contact pigs and replicated more efficiently in differentiated swine tracheal cells compared to the virus containing the wildtype human-origin HA. These results show that variants are selected quickly after replication of human-origin HA in pigs, leading to improved fitness in the swine host, likely contributing to transmission.

**Importance:** Influenza A viruses (IAV) cause respiratory disease in several species, including humans and pigs. The bidirectional transmission of IAV between humans and pigs plays a significant role in the generation of novel viral strains, greatly impacting viral epidemiology. However, little is known about the evolutionary processes that allow human IAV to become established in pigs. In this study, we generated reassortant viruses containing human seasonal HA and NA on different constellations of internal genes and tested their ability to replicate and transmit in pigs. We demonstrated that a virus containing a common internal gene constellation currently found in U.S. swine was able to transmit efficiently via the respiratory route. We identified a specific amino acid mutation that was fixed in the respiratory contact pigs that was associated with improved replication in primary swine tracheal epithelial cells, suggesting it was crucial for the transmissibility of the human virus in pigs.

## INTRODUCTION

Influenza A viruses (IAV) are enveloped viruses known to cause respiratory disease in several species, including humans and pigs. IAV has a wide host range and can adapt and cross between species (1). IAV circulates worldwide in pigs, with various strains and subtypes prevalent among swine populations. The major subtypes of IAV of swine include H1N1, H3N2, and H1N2, and considerable genetic and antigenic diversity exists within these subtypes (2).

Hemagglutinin (HA) is the major surface glycoprotein of IAV and is a critical component in determining host specificity due to its role in initial binding and entry into the host cell (1, 3, 4). The HA binds via its receptor binding site (RBS) to sialic acids (SA) in the host cell in the conformation of either SA-α-2,3 linkage or SA-α-2,6 linkage. SA-α-2,3 conformation is dominantly expressed in avian species, particularly in the respiratory and intestinal epithelial cells. SA-α-2,6 conformation is dominantly expressed in humans, usually in the upper respiratory tract (5). Thus, the host-specificity of the HA is affected by its ability to bind to these receptors (6). Specific amino acid substitutions in or near the RBS can change receptor-binding preference and facilitate host jumps (4). Specific amino acid substitutions have been identified as important in defining the receptor binding profile and to enable species crossing, such as the 226 position from H3 and H9 IAVs (7, 8).

Pigs are a unique host for IAV as they are known to express both types of SA linkages, allowing opportunity for infection with both avian and human origin IAV strains (9–11), increasing the likelihood of reassortment to occur. One example was the generation of the triple-reassortant internal gene constellation (TRIG) that emerged in the late 90s (12). The TRIG cassette contains internal gene segments derived from swine (matrix-M, non-structural-NS, and nucleoprotein-NP), human (polymerase basic 1-PB1), and avian (polymerase acidic-PA and PB2) IAVs forming a gene constellation that has remained well conserved. Previous studies have shown the TRIG cassette is prone to stably incorporate HA and NAs from various origins, potentially explaining the increased evolutionary rate of swine IAV and increased diversity after its introduction (13). The TRIG cassette also contributed to the generation of the pandemic H1N1 virus (H1N1pdm09) (14). Since then, genes of the H1N1pdm09 have been incorporated into the backbone of swine IAVs in North America (15), and most of the swine strains circulating in the United States contain a combination of internal genes of the TRIG and H1N1pdm09 lineages, with most of the genetic diversity arising from mutations in the HA and NA (16).

Frequent interspecies transmission is known to occur between humans and pigs. Numerous introductions of human IAV to pigs were identified from 1990 to 2011, including seasonal H1 and H3 viruses and continual detection of H1N1pdm09 (2, 17, 18). These human-to-swine spillover events have resulted in the establishment of many novel virus lineages circulating in pigs globally, contributing to the great genetic and antigenic diversity of swine IAV (17, 19). One such lineage resulted from a recent spillover from a human H3N2 seasonal virus that became one of the predominant H3N2 lineages in US swine and continues to evolve since its first detection in 2012 (19, 20). Although IAV are frequently exchanged between humans and pigs, most human-origin viruses reassort with swine strains after the spillover and the persisting human-origin genes show significant changes compared to the human IAV ancestors, particularly the surface genes. However, the molecular determinants enabling the adaptation of human-origin IAVs to pigs are still not clear.

Previous work has shown that wholly human H3N2 viruses do not typically replicate efficiently in pigs, and transmission is rarely observed (19, 21). Thus, to evaluate the molecular changes during replication and transmission of human-origin H3N2 surface genes in pigs, a reassortant strain was generated via reverse genetics containing human seasonal HA and NA genes in a common internal gene constellation currently found in U.S. swine. We showed this reassortant virus resulted in transmission in pigs and identified mutations in the HA gene of the resultant virus that were associated with improved transmission and replication in primary swine tracheal epithelial cells (pSTECs). Our results suggest that advantageous mutations are selected quickly in the HA gene of human seasonal IAV during replication in swine.

## MATERIALS AND METHODS

### Viruses

The wild type (wt) H3N2 human isolate A/Victoria/361/2011 (A/VIC/11) was obtained from St. Jude Children’s Research Hospital (kindly provided by Dr. Richard Webby). This virus was incorporated into the 2012-2013 human influenza vaccine for the Northern hemisphere (22). The wild type H3N1 swine isolate A/Swine/Missouri/A01410819/2014 (sw/MO/14) was obtained from the USDA National Veterinary Service Laboratories Swine IAV repository (19) and used as an example of human-origin virus adapted to swine. The swine-origin virus A/turkey/Ohio/313053/2004 (ty/OH/04), which contains the triple reassortant internal gene (TRIG) constellation, was used as the source for TRIG genes and to generate one of the control viruses for *in vitro* studies. Four viruses were generated by reverse genetics using cloned cDNAs: VIC11rg encodes all genes from A/VIC/11; VIC11p virus encodes the M gene from H1N1pdm09 and the remaining seven genes from A/VIC/11; VIC11pTRIG virus contains the HA and NA from A/VIC/11, the M from H1N1pdm09, and the remaining five genes of ty/OH/04; and ty/OH/04p virus encodes the M from H1N1pdm09 and the remaining seven genes of ty/OH/04. One mutant virus was generated: VIC11pTRIG_A138S, which encodes the same genes as VIC11pTRIG except for the A138S mutation in the HA (H3 numbering). All viruses used in this study are described in Table 1. Viruses were generated using an eight-plasmid reverse genetics system based in the bidirectional plasmid vector pDP2002, as previously described (23, 24). The mutant was generated by using the Phusion Site Directed Mutagenesis kit (ThermoFisher, Waltham, MA) with specific mutagenesis primers. Full-length sequencing of viral stocks was performed to verify gene combinations either by next generation sequencing (NGS) or Sanger sequencing. Viruses were propagated in Madin-Darby canine kidney (MDCK) cells for stock preparation.

**Table 1.** List of viruses used for this study.

| Viruses | Gene Constellation |  |  |  |  |  |  |  |
| --- | --- | --- | --- | --- | --- | --- | --- | --- |
|  | PB2 | PB1 | PA | HA | NP | NA | M | NS |
| A/VIC/11 | A/VIC/11 | A/VIC/11 | A/VIC/11 | A/VIC/11 | A/VIC/11 | A/VIC/11 | A/VIC/11 | A/VIC/11 |
| VIC11rg | A/VIC/11 | A/VIC/11 | A/VIC/11 | A/VIC/11 | A/VIC/11 | A/VIC/11 | A/VIC/11 | A/VIC/11 |
| VIC11p | A/VIC/11 | A/VIC/11 | A/VIC/11 | A/VIC/11 | A/VIC/11 | A/VIC/11 | A/CA/09 | A/VIC/11 |
| VIC11pTRIG | *ty/OH/04* | *ty/OH/04* | *ty/OH/04* | A/VIC/11 | *ty/OH/04* | A/VIC/11 | A/CA/09 | *ty/OH/04* |
| VIC11pTRIG_A138S | *ty/OH/04* | *ty/OH/04* | *ty/OH/04* | A/VIC/11* | *ty/OH/04* | A/VIC/11 | A/CA/09 | *ty/OH/04* |
| sw/MO/14 | A/CA/09 | A/CA/09 | A/CA/09 | sw/MO/14 | A/CA/09 | sw/MO/14 | A/CA/09 | A/CA/09 |
| *ty/OH/04p* | *ty/OH/04* | *ty/OH/04* | *ty/OH/04* | *ty/OH/04* | *ty/OH/04* | *ty/OH/04* | CA/09 | *ty/OH/04* |

A/VIC/11, A/Victoria/361/2011/H3N2; *ty/OH/04*, A/Turkey/Ohio/313053/2004/H3N2; A/CA/09, A/California/04/2009/H1N1; sw/MO/14, A/Swine/Missouri/2014/H3N2. \*, A/VIC/11 HA containing the A138S mutation

### Cell lines

Human embryonic kidney 293T cells (HEK293T) and MDCK cells were used for transfection of plasmids for generation of the viruses by reverse genetics. MDCK cells were also used for viral growth kinetics. Both cell lines were cultured at 37°C with 5% CO_2_ using Dulbecco’s modified Eagle’s medium (DMEM) supplemented with 10% fetal bovine serum (FBS) and 1% L-glutamine and 1% of antibiotic/antimycotics (Sigma-Aldrich, St. Louis, MO). Primary swine tracheal epithelial cells (pSTECs) were used for viral growth kinetic studies. pSTECs were harvested from trachea samples of 5 month-old pigs kindly provided by the University of Georgia, College of Agriculture, by trimming and digesting the tissue in DMEM/F12 media containing 1.5mg/ml of pronase (Sigma-Aldrich, St-Louis, MO) with antibiotic/antimycotic (Life Technologies, Waltham, MA) at 4°C for 24 hours. The harvested pSTECs were then cultured and differentiated in Air-Liquid-Interface (ALI) conditions in collagen-coated transwell inserts (Corning, New York, NY) using TEC plus and TEC plus ALI media (respectively), as previously described (25). The pSTECS were cultured for a minimum of 3 weeks at 37°C with 5% CO_2_. The basal media of the transwell inserts were replaced every 48 hours. Cilia activity was checked every 2~3 days and trans-epithelial electrical resistance (TEER) was measured to ensure confluency of the cells using the EVOM meter (World Precision Instruments, Sarasota, FL).

### In vivo study

Eighty 3-week-old cross-bred healthy pigs were obtained from a herd free of IAV and porcine reproductive and respiratory syndrome virus (PRRSV). Prior to the start of the study, pigs were treated with ceftiofur crystalline free acid (Zoetis Animal Health, Parsippany, NJ) and enrofloxacin (Elanco Animal Health, Greenfield, IN) to reduce bacterial contaminants. Animals were demonstrated to be free of other respiratory pathogens by testing bronchoalveolar lavage fluid (BALF) at the end of the study for PRRSV, porcine circovirus type 2 (PCV2), and *M. hyopneumoniae* nucleic acid by real-time RT-PCR (VetMax, Life Technologies, Carlsbad, CA) and shown to be seronegative to IAV antibodies by a commercial ELISA kit (AI MultiS-Screen kit, IDEXX, Westbrook, ME). Pigs were divided into six groups, housed in biosafety level 2 (BSL2) containment, and cared for in compliance with the Institutional Animal Care and Use Committee of the National Animal Disease Center-USDA-ARS.

Pigs (n=10/group) were inoculated intranasally (1 ml) and intratracheally (2 ml) with 10^5^ 50% tissue culture infective dose (TCID_50_) per ml of each assigned virus: A/VIC/11, VIC11rg, VIC11p, VIC11pTRIG, and sw/MO/14. Five pigs were assigned as negative controls (NC). Inoculation was performed under anesthesia, using an intramuscular injection of a cocktail of ketamine (8 mg/kg of body weight), xylazine (4 mg/kg), and Telazol (6 mg/kg) (Zoetis Animal Health, Parsippany, NJ). Five contact pigs were placed in separated raised decks in the same room as each inoculated group at 2 days post inoculation (dpi) to evaluate respiratory (airborne) transmission. Pigs were observed daily for clinical signs of respiratory disease. Nasal swabs (FLOQSwabs, Copan Diagnostics, Murrieta, CA) were collected from 0 to 5 dpi for directly inoculated pigs and from 0- to 5-, 7-, and 9-days post contact (dpc) for respiratory contacts, placed in 2 ml minimal essential medium (MEM), and frozen at −80°C until used. Two pigs died from causes unrelated to IAV, leaving eight pigs in the VIC/11wt group. Primary pigs were humanely euthanized with a lethal dose of pentobarbital (Fatal Plus, Vortech Pharmaceuticals, Dearborn, MI) and necropsied at 5 dpi. Postmortem samples included BALF, trachea and right cardiac or affected lung lobe. Indirect contact pigs were humanely euthanized at 15 dpc.

### Virus titers in nasal swabs and lungs

For virus isolation, nasal swab (NS) samples were filtered (0.45 μm) and 200 μl were plated in 24-well plates onto confluent MDCK cells washed twice with phosphate-buffered saline (PBS), as previously described (26). Tenfold serial dilutions in serum-free Opti-MEM (Gibco^®^, Life Technologies, Carlsbad, CA) supplemented with 1 μg/ml tosylsulfonyl phenylalanyl chloromethyl ketone (TPCK)-trypsin and antibiotics were made with each BALF and virus isolation-positive NS sample. Each dilution was plated in triplicate onto PBS-washed confluent MDCK cells in 96-well plates. At 48 h, plates were fixed with 4% phosphate-buffered formalin and stained using immunocytochemistry with an anti-influenza A virus nucleoprotein monoclonal antibody as previously described (27). TCID_50_/ml virus titers were calculated for each sample according to the method of Reed and Muench (28).

### Pathological examination of lungs

At necropsy, lungs were removed and evaluated for the percentage of the lung affected with purple-red consolidation typical of IAV infection. The percentage of the surface affected by pneumonia was visually estimated for each lung lobe, and a total percentage for the entire lung was calculated based on weighted proportions of each lobe to the total lung volume (29). Tissue samples from trachea and lung were fixed in 10% buffered formalin for histopathologic examination. Tissues were routinely processed and stained with hematoxylin and eosin. Microscopic lesions were evaluated by a veterinary pathologist blinded to treatment groups and scored according to previously described parameters (30). Trachea and lung tissues were assessed for IAV-specific antigen using immunohistochemistry (IHC) and scored as previously described (30). Individual scores were summed and a composite score for each pig was computed for lung and trachea microscopic lesions.

### Serology

Serum samples were collected from indirect contact pigs at 15 dpc to check for seroconversion by using hemagglutination inhibition (HI) assay. Prior to HI, sera were treated with receptor-destroying enzyme (Sigma-Aldrich, St. Louis, MO), heat inactivated at 56°C, and adsorbed with 50% turkey red blood cells (RBC). HI assays were performed with either A/VIC/11 or sw/MO/14 as antigens and 0.5% turkey RBCs using standard techniques (31). Reciprocal titers were divided by 10 and log2 transformed and reported as the geometric mean.

### Next Generation Sequencing (NGS) and Variant Analysis

Viral RNA was extracted from NS samples and tissue culture supernatants via the Magmax AI/ND extraction kit (ThermoFisher, Waltham, MA) according to the manufacturer’s instructions. After extraction, viral RNA was amplified by a one-step multi-segment RT-PCR reaction using Superscript III reverse transcriptase (Invitrogen, Carlsbad, CA) as previously described (32). PCR conditions are 55°C for 2 min, 94°C for 2 min, followed by 35 cycles at 94°C for 30 sec, 50°C for 30 sec, 68°C for 3 min, and final elongation of 4 min at 68°C. Influenza whole genome sequencing libraries were prepared using the Nextera XT DNA library preparation kit (Illumina, San Diego, CA). Libraries were selected and purified using 0.7× Agencourt AMPure XP Magnetic Beads (Beckman Coulter Life Sciences, Indianapolis, IN, USA) and samples were normalized to 4 nM and pooled. Fragment size distribution was analyzed using the High sensitivity DNA kit (Agilent, Santa Clara, CA, USA). Pooled libraries were sequenced via the high-throughput MiSeq platform (Illumina, San Diego, CA) using the 300-cycle MiSeq Reagent Kit in a paired end format (Illumina, San Diego, CA, USA). Full genome assembly was performed using a pipeline from previously published methods (32). Variant calling was performed using LoFreq 2.1.3.1(33) with a frequency threshold of 0.02 based on replicate sequence runs for the same control samples, a minimum depth of coverage of 100 and a central base quality score of Q30 or higher. Quantification of viral diversity within and between hosts were calculated using the consensus sequences and single nucleotide variants within all hosts derived from the variant call data utilizing the Lasergene bioinformatics package (DNASTAR, Madison, WI).

### Mapping of variants on the HA protein

Three-dimensional (3D) analysis was conducted on the HA protein of the VIC11pTRIG variants identified in the study. Protein sequences were checked using DNAStar (Madison, WI, USA) and ExPASy (SIB bioinformatics resources), submitted to the I-TASSER (University of Michigan, Ann Arbor, MI, USA), and visualized with CHIMERA 1.141 (University of California, San Francisco, CA, USA). Only protein structures with a positive C-score (Confidence score) were selected for analysis.

### Viral Growth Kinetics

Viral growth kinetics were performed for strains VIC11pTRIG, VIC11pTRIG_A138S and ty/OH/04p in MDCKs and pSTECs. Both cells were infected at a multiplicity of infection (MOI) of 0.05. Infection media for MDCK consisted of Opti-MEM (Thermo Fisher Scientific, Waltham, MA, USA) containing 1 ug/ml TPCK trypsin (Worthington Biochemicals, Lakewood, NJ, USA) with 1% of antibiotic/antimycotic solution (Sigma-Aldrich St. Louis, MO, USA). Infectious media for pSTECs consisted of F12/DMEM media supplemented with 0.075% BSA (Sigma-Aldrich St. Louis, MO, USA) and 200nm GlutaMax (Thermo Fisher Scientific, Waltham, MA, USA). Cells were infected in triplicates and supernatant samples were collected at 0, 12, 24, 48 and 72 hours post infection (hpi). Virus titers were quantified and titrated by real-time PCR (QuantaBio ToughMix®, VWR International, Radner, PA) with a TCID_50_/ml equivalent as previously described (34). Results from two independent experiments were compiled for analysis.

### Statistical analysis

All statistical analyses were conducted by using GraphPad Prism version 9.3.1 (GraphPad Software, San Diego, CA) with p≤0.05 considered significant. Statistical methods include nonparametric one-way ANOVA, two-way ANOVA, and nonparametric t-tests.

## RESULTS

### The reassortant VIC11pTRIG replicated and caused disease in pigs

Pigs (n=10/virus) were challenged with A/VIC/11, VIC11rg, VIC11p, VIC11pTRIG, and sw/MO/14. BALF were collected from directly infected pigs at 5 dpi, when the percentage of lung lesions was assessed (Fig. 1A-B). Confirming our previous results (19), viral titers were only detected in the BALF of 2 pigs infected with the A/VIC/11 and VIC11rg wholly human seasonal viruses, and from only 1 pig infected with the VIC11p. In contrast, 7 pigs from the VIC11pTRIG infected group had detectable virus titers in the BALF, while virus was detected in all pigs from the sw/MO/14 infected group (Fig. 1A). Pigs infected with the sw/MO/14 showed significantly higher titers compared with all other groups. Although replication was confirmed in a few pigs infected with A/VIC/11, VIC11rg, or VIC11p, titers were low (below 10^2.5^ TCID_50_/ml), bringing the group average to below the limit of detection (Fig. 1A). Similarly, IAV-specific antigen staining was detected by IHC in sw/MO/14-challenged pigs, but signals were not observed in any of the other challenged pigs (Table 2).

**Figure 1.**
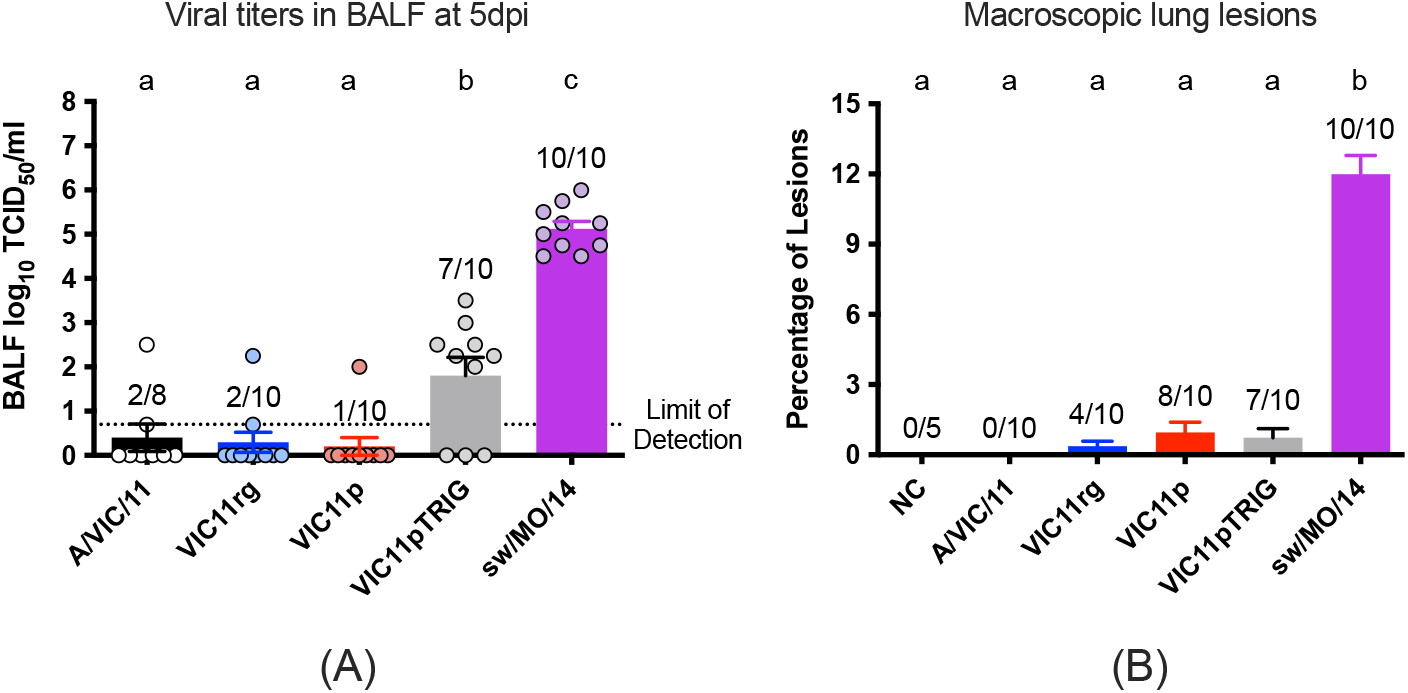
Viral titers in the lungs and macroscopic lung lesions of pigs directly infected with reassortant viruses. (A) Viral titers in bronchoalveolar lavage fluid (BALF) collected at 5 days post infection (dpi) from pigs directly infected with A/VIC/11, VIC11rg, VIC11p, VIC11pTRIG, or sw/MO/14. Values are shown as mean TCID_50_/ml titers ± standard error of the mean, with scattered dots representing individual pigs. (B) Percentage of the lungs affected with purple-red consolidation at 5 dpi. Numbers indicated above the error bar depicts the number of positive or affected pigs/total number of pigs in the group. Different lowercase letters indicate significant difference at *P<0.05*.

**Table 2.** Lung and trachea microscopic pathology and virus detection in pigs challenged with reassortant viruses and negative controls^a^.

| **Virus** | **Microscopic lung lesions score<sup>b</sup>** | **Lung IHC score<sup>c</sup>** | **Microscopic trachea lesions score<sup>c</sup>** | **Trachea IHC score<sup>d</sup>** |
| --- | --- | --- | --- | --- |
| **NC** | 0.25 ± 0.16<sup>A</sup> | 0.0 ± 0.0<sup>A</sup> | 0.20 ± 0.20<sup>A</sup> | 0.0 ± 0.0<sup>A</sup> |
| **A/VIC/11** | 0.78 ± 0.19<sup>A</sup> | 0.0 ± 0.0<sup>A</sup> | 0.75 ± 0.22<sup>A</sup> | 0.0 ± 0.0<sup>A</sup> |
| **VIC11rg** | 0.60 ± 0.18<sup>A</sup> | 0.0 ± 0.0<sup>A</sup> | 0.20 ± 0.15<sup>A</sup> | 0.0 ± 0.0<sup>A</sup> |
| **VIC11p** | 0.58 ± 0.15<sup>A</sup> | 0.0 ± 0.0<sup>A</sup> | 0.05 ± 0.05<sup>A</sup> | 0.0 ± 0.0<sup>A</sup> |
| **VIC11pTRIG** | 0.48 ± 0.21<sup>A</sup> | 0.0 ± 0.0<sup>A</sup> | 0.65 ± 0.18<sup>A</sup> | 0.02 ± 0.02<sup>A</sup> |
| **sw/MO/14** | 9.43 ± 0.61<sup>B</sup> | 5.3 ± 0.4<sup>B</sup> | 2.13 ± 0.29<sup>B</sup> | 2.58 ± 0.23<sup>B</sup> |

<sup>a</sup> Results are shown as means ± standard errors of the means. Different uppercase letters within the same column indicate significant difference at P<0.05. <sup>b</sup> The range of possible scores is 1 to 22. <sup>c</sup> The range of possible scores is 1 to 8. <sup>d</sup> The range of possible scores is 1 to 4. IHC, immunohistochemistry.

As expected, sw/MO/14 induced a high percentage of pneumonia in all pigs, significantly higher compared to all other groups (Fig. 1B). The average percentage of macroscopic lesions were very low in the groups challenged with A/VIC/11, VIC11rg, VIC11p, VIC11pTRIG, with no significant difference compared with the negative (non-infected) group. Similarly, pigs infected with sw/MO/14 showed the highest average microscopic lung and trachea lesion scores (Table 2), significantly higher than the other groups.

Although some pigs infected with A/VIC/11 and VIC11pTRIG showed slightly higher trachea lesion scores compared to the other groups challenged with viruses containing the A/VIC/11 HA (Table 2), no significant differences were observed among these groups nor compared to the negative controls.

### The reassortant VIC11pTRIG transmitted between pigs

Nasal swab viral titers were assessed in directly inoculated pigs from 1 to 5 dpi and in respiratory contacts from 1 to 5, 7, and 9 dpc. Consistent with replication in the lungs, only a small number of pigs directly infected with the A/VIC/11 or VIC11rg shed low virus titers, with only 3 and 2 pigs, respectively, testing positive by virus isolation at any time-point (Fig. 2A). Compared to pigs infected with wholly human seasonal virus (A/VIC/11 or VIC11rg), more pigs directly infected with VIC11p shed virus, with almost all pigs (8 out of 10) being positive at least in one time-point. However, no statistical difference was observed compared to the A/VIC/11 or VIC11rg groups. In contrast, virus was detected in the nasal swabs of all pigs infected with VIC11pTRIG and sw/MO/14 at all time-points, and viral titers were significantly higher in comparison with the other three groups for most time-points (Fig. 2A). Consistent with little to no shedding observed in A/VIC/11-, VIC11rg-, and VIC11p-directly infected pigs, none of the respiratory contact pigs (n=5/group) that were in the same room as these groups were positive at any time-point (Fig. 2B). In contrast, all pigs in contact with pigs infected with VIC11pTRIG or sw/MO/14 were positive in at least one time-point, starting mostly at 4dpc (Fig. 2B). Only one pig in the VIC11pTRIG contact group was positive at 3 dpc (data not shown). Consistent with these results, all pigs in these two groups were seropositive by hemagglutination inhibition (HI) assay at 15 dpc (Fig. 2C).

**Figure 2.**
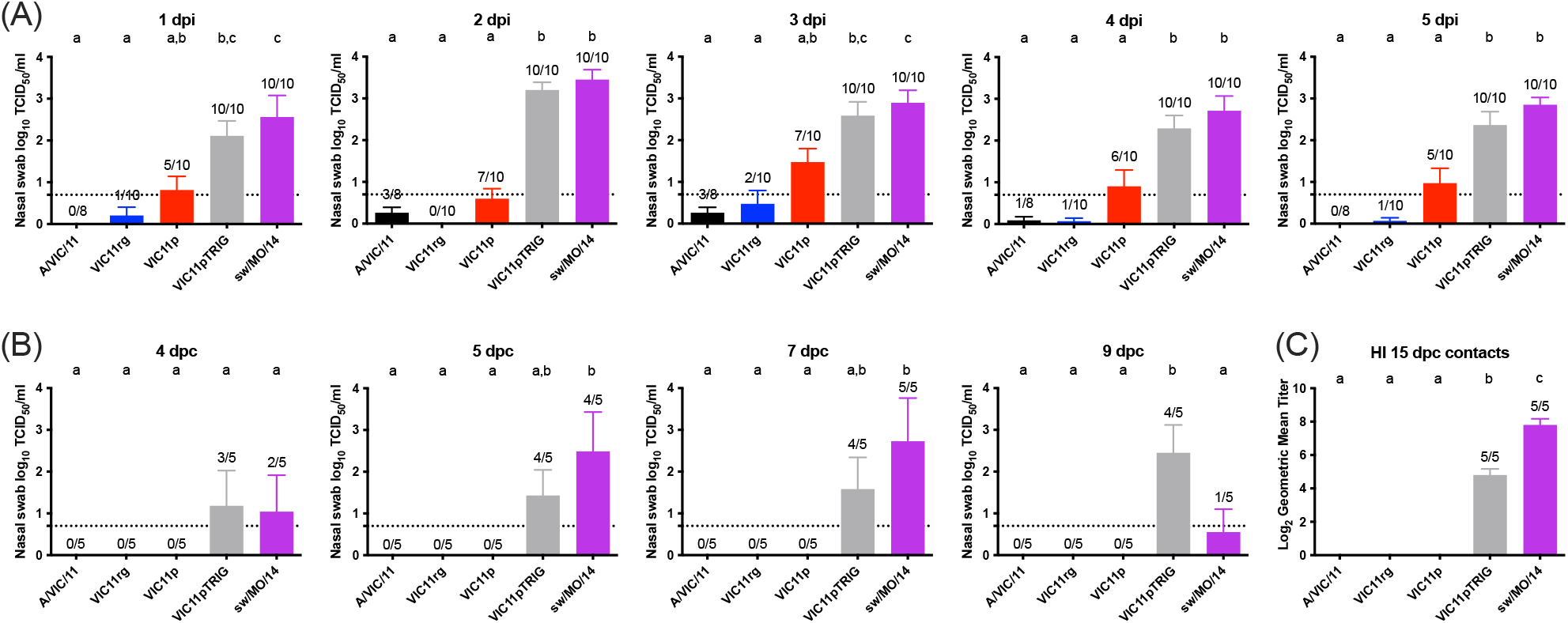
Viral titers in nasal swabs of directly infected and respiratory contact pigs after infection with human-origin reassortant viruses and seroconversion of respiratory contact pigs. Nasal swabs samples were collected from (A) directly infected pigs daily from 1 to 5 days post infection (dpi) and from (B) respiratory contact pigs daily from 1 to 5, then 7- and 9-days post contact (dpc). (C) Antibody response in respiratory contact pigs at 15 dpc measured by HI assay. Groups were infected with A/VIC/11, VIC11rg, VIC11p, VIC11pTRIG, or sw/MO/14. Values are shown as mean TCID_50_/ml titers or geometric mean titers ± standard error of the mean. Numbers indicated above the error bar depicts the number of positive pigs/total number of pigs in the group. Different lowercase letters indicate significant difference at *P<0.05*.

### Minor variants are selected quickly after replication of the reassortant VIC11pTRIG in pigs

Nasal swab (NS) samples from 5 directly inoculated pigs collected at 1, 3, and 5 dpi, and 5 respiratory contacts collected at 5, 7 and 9 dpc in the VIC11pTRIG group were selected for sequencing by NGS. Samples from directly infected pigs with highest TCID_50_/ml titers in any given time-point (>10^2.75^ TCID_50_/ml) were selected for sequencing, and all respiratory contacts. Of the 30 NS samples selected for NGS sequencing, 9 samples from directly infected pigs (pigs 375, 378, 340 in all selected time-points) and 12 samples from respiratory contact pigs (pigs 407, 408, 409, 410 in all selected time-points) were amplified by MS-RT-PCR, and 18 produced complete genome assemblies by NGS. Viral genomes were analyzed for HA and NA variants. A total of 3 dominant variants were discovered in the HA: A138S, V186G, and F193Y. A138S emerged in two directly infected animals, becoming dominant (77% of viruses) in one animal by 5 dpi (Fig. 3A). A138S was fixed (99% of viruses) in all 4 contact pigs soon after transmission (Fig. 3B). The V186G mutation was present in 2.6% of viruses in one directly infected animal at 5 dpi, was not detectable in the contact pigs soon after transmission but became dominant in one animal on 9 dpc with a frequency of 60% (Fig. 3C-D). The F193Y mutation was only detected in one directly inoculated pig at a frequency of 56% by 3 dpi, declining on 5 dpi to 27% (Fig. 6E). F193Y was transmitted to one pig with a frequency of 9% at 5 dpc and at a frequency of 4% in another animal by 9 dpc (Fig. 6F). In addition to the major variants, several low-frequency variants were identified along the HA, in a total of 46 positions (Table 3). No dominant variants were observed in the NA or other viral gene segment.

**Figure 3.**
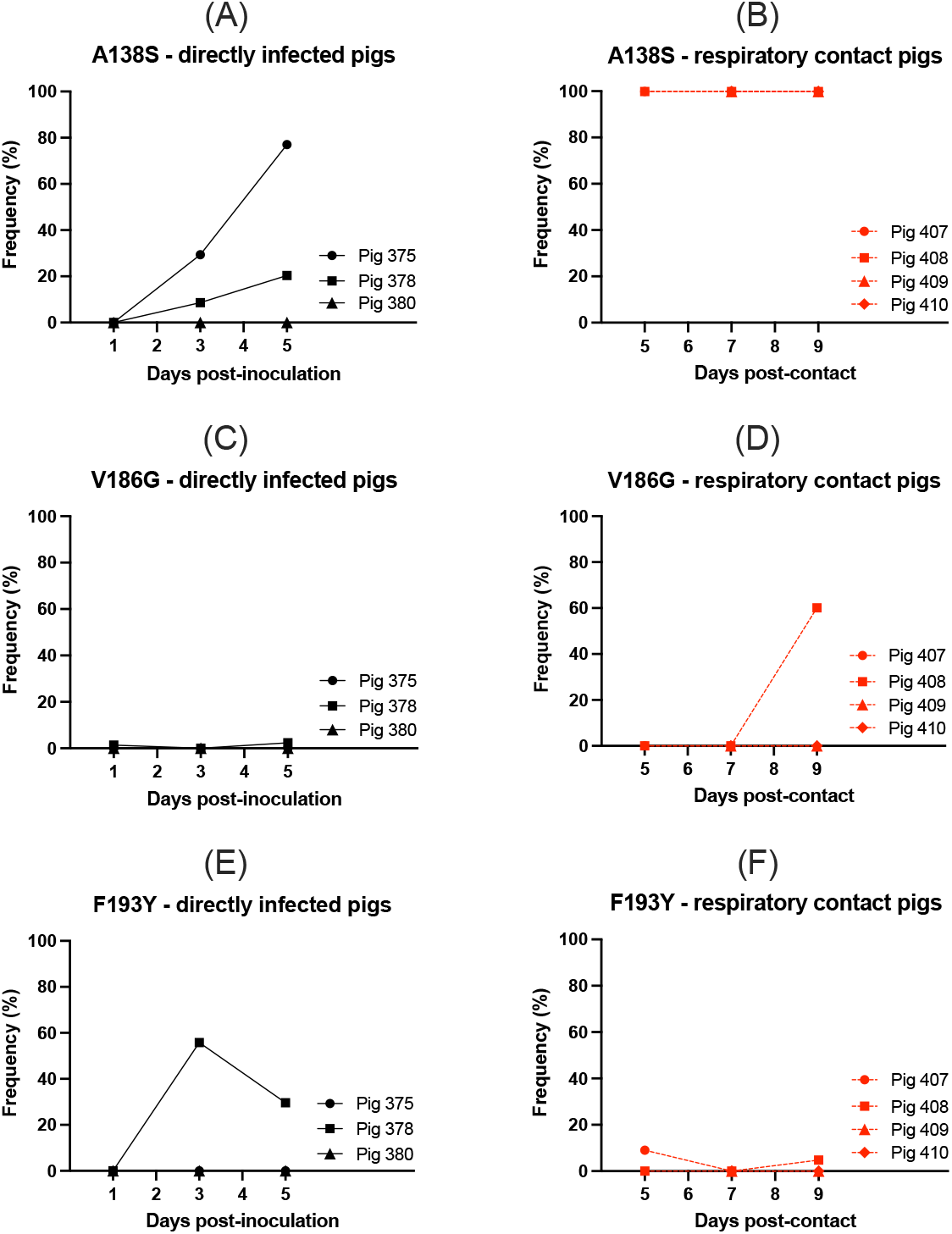
Variants identified in directly infected and respiratory contact pigs in the group infected with the VIC11pTRIG reassortant. Dominant variants detected in each directly infected and respiratory contact pigs are shown. Variants with a frequency higher than 50% that were detected in nasal swabs in at least one occasion were considered dominant. (A,B) depicts mutation A138S; (C,D) depicts mutation V186G; (E,F) depicts mutation F193Y. All positions are based on H3 numbering.

**Table 3.**
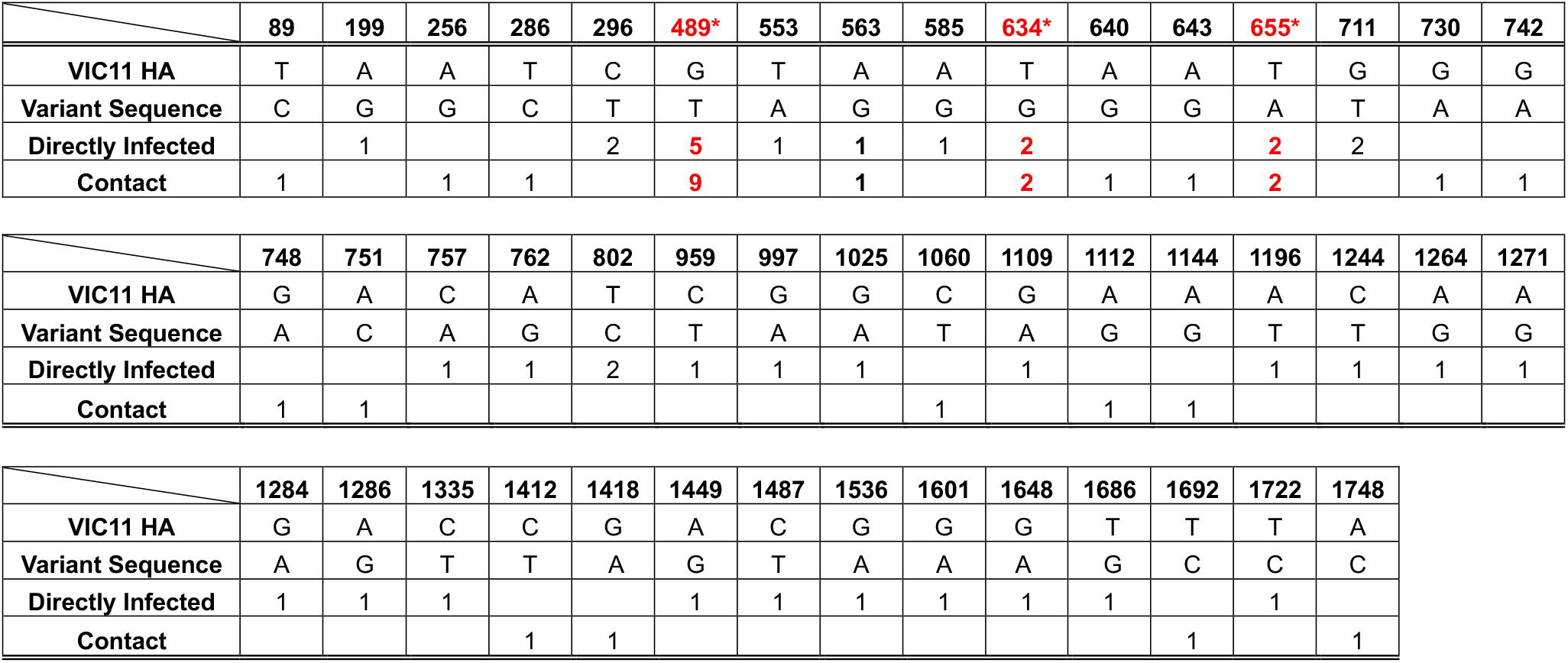
Variants identified in directly infected and respiratory contact pigs infected with VIC11pTRIG. List of variants in nasal swab samples from directly infected and respiratory contact pigs in this study. The numbers on the top represent the nucleotide positions. The VIC11 HA represents the HA sequence of A/Victoria/361/2011/H3N2, which is the same as VIC11pTRIG. Substitutions at the positions relative to the reference sequence present in ≥2% of variant viruses are shown. Number of samples containing each variant for either directly infected or respiratory contacts are listed. *Positions 489, 634, 655 correspond to amino acid residues 138, 186, 193, respectively.

We plotted the location of the 3 mutations on the HA head and found all residues were located close to H3 antigenic and binding sites (Fig. 4). The A138S mutation is located under the 220 loop which is one of the major structures forming the receptor binding site (RBS). The F193Y and V186G are located next to the 190 loop, another structure that forms the RBS.

**Figure 4.**
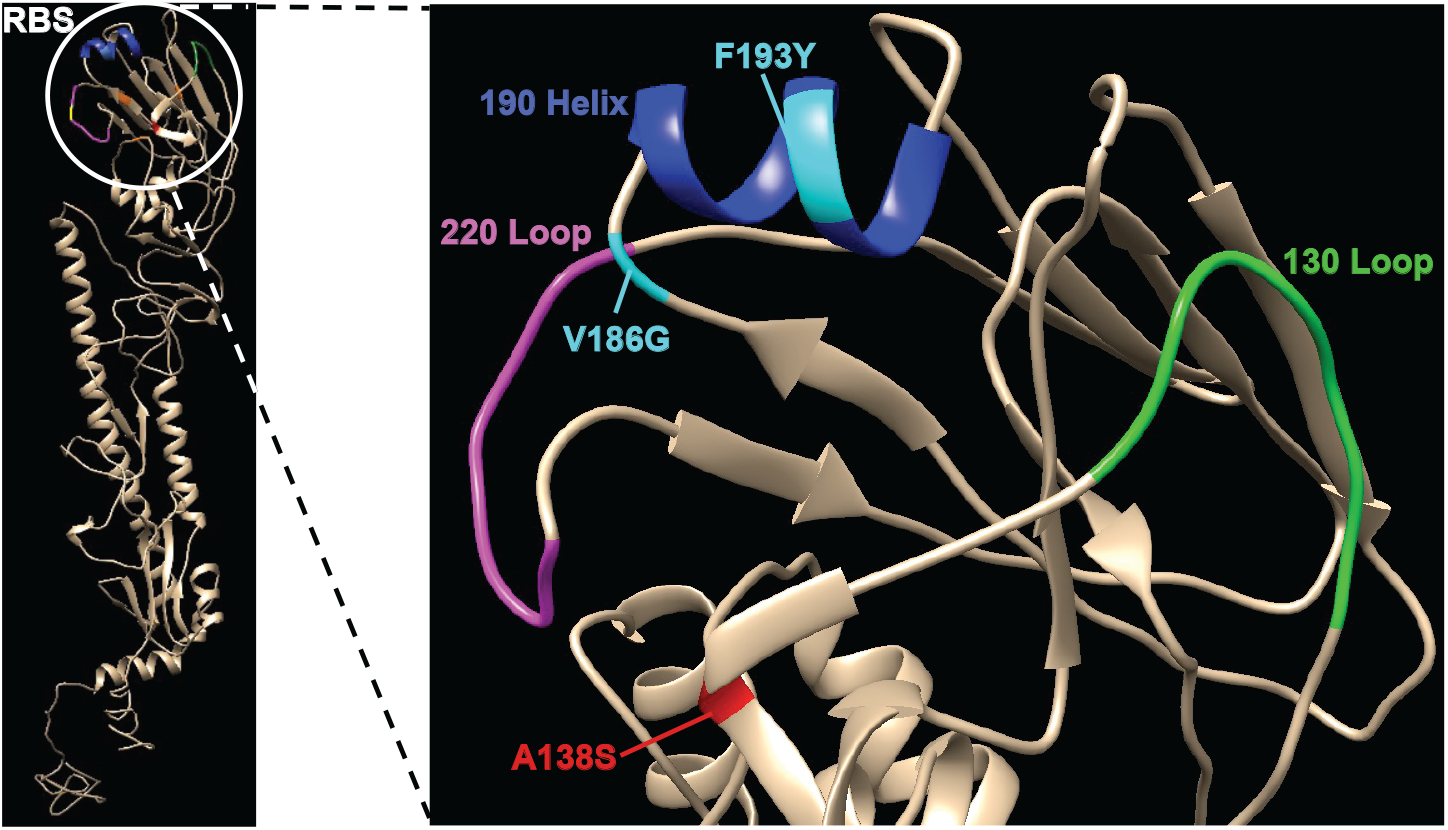
Location of mutations identified on the Vic11pTRIG HA. Dominant variants identified in this study on the HA of the Vic11pTRIG are highlighted on the structure of the human A/Victoria/361/2011 H3. Red, residue A138S; cyan, residues V186G and F193Y. The 190 helix, 130 loop, and 220 loop of the receptor binding site (RBS) are highlighted.

### Mutation A138S improves replication of the human seasonal HA of VIC11pTRIG in swine cells

Viral growth kinetics were evaluated in pSTECs and MDCKs to compare the replication of the A138S variant (VIC11pTRIG_A138S) in comparison to VIC11pTRIG (Fig. 5). This variant was selected because it was the only variant that was transmitted and became dominant/fixed in all contact pigs. ty/OH/04p was used as control and contains the same backbone as the other two viruses. The VIC11pTRIG_A138S showed more efficient replication in pSTECs at all time-points (Fig. 5A) in comparison to VIC11pTRIG containing the human seasonal HA, with the highest difference at 72 hpi. Replication of VIC11pTRIG_A138S was similar to ty/OH/04p up to 24 hpi but ty/OH/04p showed significantly higher titers at 48 and 72 hpi. No significant difference in replication kinetics was observed in MDCK cells for any of the viruses (Fig. 5B).

**Figure 5.**
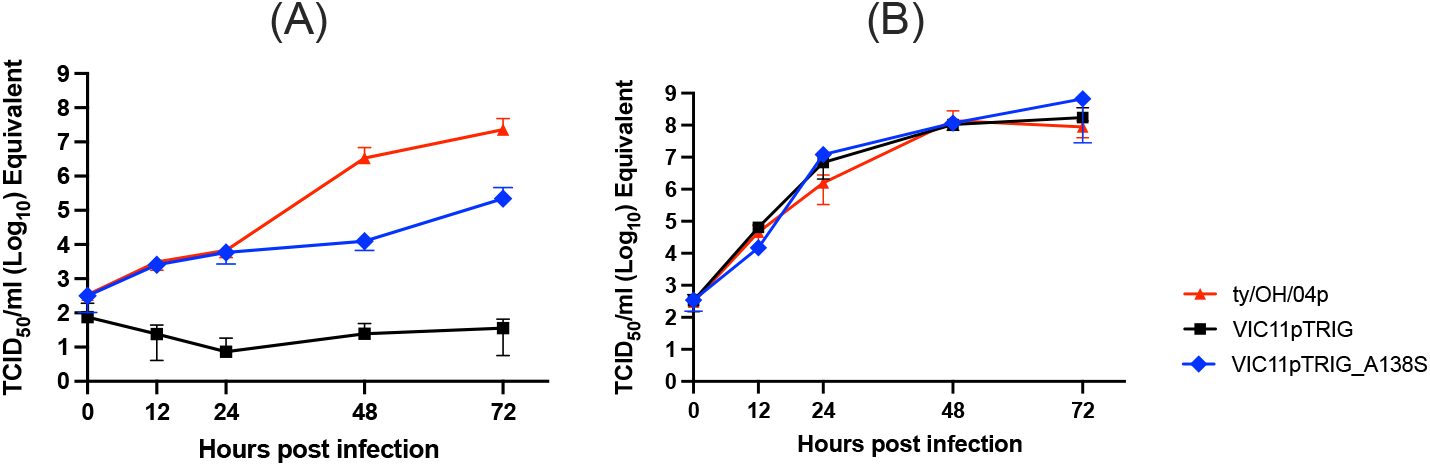
*In vitro* replication kinetics of mutant containing A138S substitution in primary swine tracheal epithelial cells (pSTECs) and MDCK cells. Viral growth kinetics of A138S mutant virus (VIC11pTRIG_ A138S), VIC11pTRIG reassortant and ty/OH/04p in (A) pSTECs and (B) MDCK cells. pSTECs and MDCK cells were infected at an MOI of 0.5, and supernatants were collected at 0, 12, 24, 48, and 72 hours post infection (hpi). Viral titers were quantified by real-time PCR with a TCID_50_/ml equivalent. Values are shown as mean TCID_50_/ml titers ±standard error of the mean.

## DISCUSSION

Determining the evolutionary processes of influenza virus cross-species transmission is key to understanding the mechanisms that control the emergence of new influenza viruses and establishment in a new host population. Spillover events of IAV between humans and swine are common and contribute to the extensive diversity of IAV circulating in pigs. The TRIG backbone seems to have an important part in expanding this diversity as it has the capability to incorporate HA and NA segments without destabilizing the overall fitness of the strains (2, 4, 19, 20). While acquiring an efficient and permissive internal gene constellation seems to be a crucial step in the adaptation of IAV to a new species, the molecular mechanisms that drive viral evolution after transmission of human-origin viruses in swine is not well understood. Thus, we generated a novel reassortant virus (VIC11pTRIG) that contains human seasonal HA and NA in a backbone highly adapted to swine and assessed its evolution during replication and transmission in pigs. Our results show that this ideal backbone significantly increases fitness of human seasonal H3N2 in pigs and is crucial for the transmissibility in pigs, allowing for the further evolution and selection of HA variants.

In this study, we have shown that acquiring the ideal combination of genes is likely the first step for the adaptation of human-origin IAV to successfully replicate and transmit in pigs. The VIC11pTRIG group showed significantly higher viral titers in nasal swabs and BALF compared to groups with full constellation of human-origin internal genes or with the addition of the H1N1pdm09 M gene. More importantly, the VIC11pTRIG was the only VIC11 reassortant able to transmit among pigs. The fitness advantage of the TRIG cassette and the importance of an ideal gene constellation had been confirmed previously in co-infection studies, in which only an H3N2 virus with a particular constellation containing the TRIG cassette was able to transmit between pigs (12). Here, the addition of the H1N1pdm09 M gene conferred a replication advantage with the human-origin internal gene constellation as a higher number of infected pigs in this group shed higher viral titers compared to other groups with human seasonal internal genes (A/VIC/11 and VIC11rg), although differences were not statistically significant. However, though directly infected pigs shed virus, there was no transmission. Interestingly, while transmission was confirmed in the VIC11pTRIG reassortant group to similar levels as in the swine-adapted sw/MO/14 group, infection resulted in little to no lung lesions and significantly lower titers in the lungs compared to the swine-adapted virus. These findings are consistent with previous studies that demonstrated reassortant viruses carrying the A/VIC/11 HA showed replication in the upper respiratory tract and limited replication in the lungs (19).

The HA is a major factor for host specificity (35, 36) and is a prime target for evolution as a human seasonal virus adapts to the swine host. Specific amino acid substitutions at or near the RBS have been shown to alter receptor binding or antigenicity of influenza viruses (5, 37, 38), which likely affects host specificity. Hence, it is reasonable to infer that once the virus acquires a gene constellation that allows effective replication, in this case the pTRIG backbone, mutations that favor receptor binding in the new host may be an additional step to an efficient transmission. Here, the 3 dominant mutations that arose, A138S, V186G, and F193Y, were all located within the antigenic site B of H3N2 (39), near or within the RBS. Thus, it is possible that these mutations, particularly the A138S that was fixed in respiratory contact animals soon after transmission, may have altered the conformation of the HA and its interaction with the host cell. Interestingly, the 138 position was shown previously to affect infectivity (40–42). The advantageous effect of the A138S mutation for replication in swine was confirmed in growth kinetic assays in pSTECs, a system that mimics the swine respiratory tract, which suggests that this mutation was crucial for the transmissibility of the VIC11pTRIG reassortant virus in our study. The V186G mutation has been detected in surveillance studies as a variant of A/VIC/11-like strains (43), and was associated with replication and evolution in an immunocompromised human (44). Although this variant was not transmitted as a major variant in pigs, it subsequently became dominant in a contact animal and it could represent one of many different evolutionary pathways that later could be selected in swine as an advantageous variant, as previously proposed for transmissibility of avian H1N1 in mammals (45). The F193Y mutation was dominant in one directly infected pig and was potentially transmitted in low frequency. However, it did not become dominant in contact pigs, suggesting this evolutionary pathway did not present an advantage in the swine host.

Although the reassortant virus that we studied here contains human seasonal HA and NA, the rest of the genome was formed by well-adapted swine IAV internal genes, limiting our ability to evaluate evolution outside of the surface genes. Nevertheless, using an artificial virus with a background known to be effective in pigs allowed us to examine the immediate virus evolution after initial replication in the swine host. As expected, the diversity in the HA was significantly higher after replication in pigs compared to other segments, including the NA, and no mutations were fixed in any other segments (data not shown). However, it is unknown whether the A138S HA mutation would have become fixed if a wholly human virus was tested.

Overall, this study suggests that for a human-origin virus to become adapted to pigs it needs an efficient internal gene constellation capable of enhancing replication, allowing for the HA diversification and selection of advantageous variants that will be transmitted between pigs. However, our study is limited to a single transmission event and further studies are needed to accurately confirm these evolutionary processes and the molecular mechanisms required for the adaptation of human-origin viruses in swine.

## ACKNOWLEDGMENTS

We thank producers, swine veterinarians, and diagnostic laboratories for participating in the USDA Swine Influenza A Virus Surveillance System. We thank Michelle Harland and Gwen Nordholm for technical assistance with laboratory techniques and Jason Huegel, Ty Standley, and Jason Crabtree for animal care.

This study was supported by the University of Georgia Office of Research, USDA-ARS, USDA-APHIS, and by an NIH-National Institute of Allergy and Infectious Diseases (NIAID) interagency agreement associated with Center of Research in Influenza Pathogenesis, an NIAID funded Center of Excellence in Influenza Research and Surveillance (HHSN272201400008C). This study was supported in part by resources and technical expertise from the Georgia Advanced Computing Resource Center, a partnership between the University of Georgia’s Office of the Vice President for Research and Office of the Vice President for Information Technology. EJA was supported in part by an appointment to the ARS, USDA, Research Participation Program, administered by the Oak Ridge Institute for Science and Education (ORISE) through an interagency agreement between the U.S. Department of Energy (DOE) and USDA. ORISE is managed by ORAU under DOE contract number DEAC05-06OR23100. Mention of trade names or commercial products in this article is solely for the purpose of providing specific information and does not imply recommendation or endorsement by UGA, USDA, DOE, or ORISE. USDA is an equal opportunity provider and employer.

